# Systems Biology Approach of Understanding Insulin Resistance: Linkage between Type 2 Diabetes & Alzheimer’s Disease

**DOI:** 10.1101/2022.04.13.463890

**Authors:** Osbert Bryan Villasis, Daniel Stanley Tan, Eduardo Mendoza, Angelyn Lao

## Abstract

Insulin resistance (IR) is a physiological condition in which cells in the body become resistant to insulin. It is a known risk factor associated to type 2 diabetes (T2D). Recently, the idea that IR plays an important role in the progression of Alzheimer’s disease (AD) has been gaining a lot of attention. Comparing the components of the insulin signalling pathway in relation to T2D and AD, there seems to be a lot of commonality. However, on what role IR plays in linking T2D and AD remains unknown. Through systems biology approach, we extended an existing mathematical model (i.e. ODE based) to study and understand the role IR plays in linking T2D and AD. The simulations, together with the experimental data collected from the literature, show that the common components in T2D and AD express the same dynamical behaviors. This result provides the bases for further modelling of the insulin signaling pathway in determining the link between T2D and AD.

## INTRODUCTION

It was the groundbreaking Rotterdam study (Ott et al., 1998) that first links type 2 diabetes (T2D) and Alzheimer’s disease (AD). The study suggested that patients with T2D increased their risk of developing dementia and AD. As the rates of the aging population increase, this landmark study has been getting more attention. Although a number of studies has provided further direct evidence to solidify the connection between T2D and AD (Han & Li, 2010; Kroner, 2009; Ott et al., 1998), but still, diabetes and AD are connected in ways that aren’t completely understood.

AD is a neurodegenerative disorder characterized by the neuropathologic findings of intracellular neurofibrillary tangles (NFT) and extracellular amyloid plaques (Aβ) that accumulate in specific brain regions (Sennvik et al., 2000). Aβ plaque formation is one of the most studied pathological hallmarks of Alzheimer’s disease (LaFerla et al., 2007; Soriano et al., 2001). Many studies have focused on either removing Aβ plaques (Mawuenyega et al., 2010; Pahnke et al., 2009; Selkoe, 2001) or interrupting the APP cleaving process (Moss et al., 2011; Loane et al., 2009; Lourenço et al., 2009).

Loss of insulin signaling in diabetes can occur by either type 1 or type 2 processes. T2D is the most common form of diabetes (Alex et al., n.d.). Unlike type 1 diabetes, patients with T2D make insulin, but their tissues are unresponsive to its effects (Kroner, 2009). This is called insulin resistance. When there isn’t enough insulin or the insulin is not used as it should be, glucose can’t get into the body’s cells. When glucose builds up in the blood instead of going into cells, the body’s cells are not able to function properly. Consequently, the process impairs the insulin signaling.

Insulin is the major hormone controlling critical energy functions such as glucose and lipid metabolism. Insulin activates the insulin receptor (IR), which phosphorylates and recruits different substrate adaptors such as the IRS family of proteins. Tyrosine phosphorylated IRS then displays binding sites for numerous signaling partners. Among them, PI3K has a major role in insulin function, mainly via the activation of the Akt/PKB(protein kinase B) and the PKCζ cascades. Activated Akt induces (1) glycogen synthesis through inhibition of GSK-3; (2) protein synthesis via mTOR(mammalian target of rapamycin) and downstream elements; and (3) cell survival through inhibition of several pro-apoptotic agents (Bad, Forkhead family transcription factors, GSK-3). Insulin stimulates glucose uptake in muscle and adipocytes via translocation of GLUT4 vesicles to the plasma membrane. GLUT4 translocation involves the PI3K/Akt pathway.

Insulin signaling also has growth and mitogenic effects, which are mostly mediated by the Akt cascade as well as by activation of the Ras/ MAPK pathway. In addition, insulin signaling inhibits gluconeogenesis in the liver, through disruption of CREB/CBP/Torc2 binding. Insulin signaling also promotes fatty acid synthesis through activation of SREBP-1C, USF1, and LXR. A negative feedback signal, emanating from Akt/PKB, PKCζ, p70 S6K, and the MAPK cascades, results in serine phosphorylation and inactivation of IRS signaling.

In Hou and colleagues’ (Hou et al., 2004) paper, they discussed about how GLUT4 regulates among the trans-Golgi network (TGN), endosomes, and plasma membrane in the absence of insulin. Interestingly, these are precisely the three main compartments in the processing of amyloid precursor protein (or APP processing) in Alzheimer’s disease. However, unlike in the muscle cells, where glucose enters through GLUT4 receptors via facilitated diffusion(Leto & Saltiel, 2012), in the brain, glucose enters passively.

Han et al. (2010) elaborated (n.d.) how insulin resistance and inflammation can be depicted as the mechanistic links between AD and T2D. As shown in Fig. 1A, Insulin resistance in peripheral tissues and organs, when coupled with relative insulin deficiency, causes T2D. On the other hand, central insulin resistance, together with reduced brain insulin levels that might have resulted from T2D, leads to accumulation of β-amyloid and, consequently, AD. As for inflammation (as shown in Fig. 1B), through its influence on islet function and peripheral insulin sensitivity, inflammation accelerates the development of T2D. Cerebrovascular and central inflammation, along with increased accumulation of β-amyloid, disrupts normal synaptic function, a starting point of AD pathological progression.

**Fig. 1.**
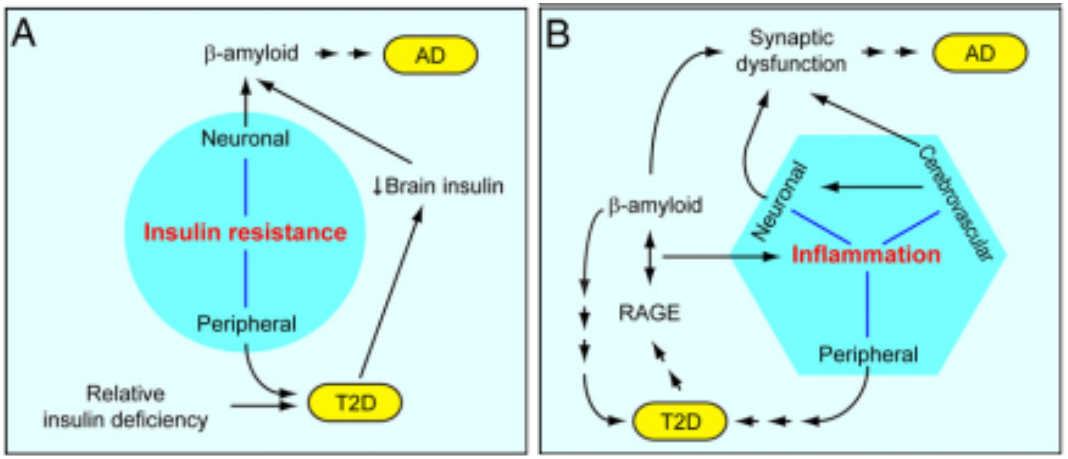
The underlying links between AD and T2D (Han & Li, 2010).

Mitochondrial dysfunction, oxidative stress and dysregulated calcium homeostasis are known to be associated with diabetes, which might also be contributory factors to the development of AD (Sung et al., 2011; Sims-Robinson et al., 2010). Oxidative stress combined with an increase in intracellular calcium result in a feedforward cycle of continued mitochondrial damage that can cause neuronal death and, hence contribute to AD pathology. During hyperinsulinemia, insulin and Aβ compete for insulin-degrading enzyme, leading to Aβ accumulation and plaque formation. And as the insulin receptor decrease, it leads to inhibition of Akt and dephosphorylation (activation) of GSK-3β (glycogen synthase kinase 3β), and results in tau hyperphosphorylation. In their paper, they emphasized that in type 2 diabetes, insulin resistance leads to both Aβ plaque formation and tau hyperphosphorylation.

Studies have suggested that T2D poses a greater risk for AD development (Aulston et al., 2013; Peila et al., 2002; Alan J. Sinclair, 2000; Arnold et al., 2018). In this study, we use systems biology approach to understand the link between Ad and T2D through insulin resistance. Systems biology is a holistic interdisciplinary approach to translate and decode complex biological systems. It is helpful in designing predictive, multiscale models that enables researchers to discover new biomarkers for disease, stratify patients based on unique profiles, and target drugs and other treatments (*What Is Systems Biology* · *Institute for Systems Biology*, n.d.; Wolkenhauer et al., 2009). Through systems biology approach, we want to better understand how AD occurs and transforms. A better understanding of the disease could help uncover ways to prevent it, delay its progression or treat the disease.

## MODELS

We first tried to find the connection or relationship of AD to other diseases, based on their shared pathways, we tried to cluster and visualize relationships with respect to shared entities on the pathways of human diseases using the Self-Organizing Map (SOM) (Sarmiento et al., 2017), which is a classical statistical cluster analysis tool. SOM is a type of artificial neural network that employs unsupervised learning that is capable of discovering patterns in datasets by reducing multi-dimensional data to a low-dimensional representation. We performed similar analysis using other clustering and similarity analysis tool (Mendoza et al., 2018). By doing so, it could provide new insights into disease classification.

Moreover, there are already various studies on the relationships between human diseases, the relationships and interactions among human genes and proteins, and the associations between genes and diseases. So, another aspect is to automate corpus annotation of biomedical journals through machine learning approach to automate the extraction of protein, genes, and diseases relationships (Laron, Andrew et al., 2017). This is helpful and useful for predicting novel associations between biomedical entities and it can also use to suggest new topics for experiments and new insights in drug design. Using the extracted and annotated corpus, through support vector machines approach, we were able to generate graph visualization of associations, with the nodes representing the diseases, while the edges representing the associations between diseases, and the thickness of an edge representing the strength of the association between the diseases which it connected, as determined by the system. We have shown that insulin resistance or dysfunction of insulin signaling, a universal feature of T2D, is the main culprit of AD progression linking T2D with AD.

In this study, the experimental data that will be the basis for model simulation are from Bränmark et al. on Insulin Signaling in Type 2 Diabetes (Brännmark et al., 2013) and Talbot et al. on brain insulin resistance in Alzheimer’s disease patients (Talbot et al., 2012). Both are focused on insulin signaling. However due to different objectives of their study, their models highlight different components of the signaling pathways. In Bränmark et al. (Brännmark et al., 2013), they included markers connected to diabetes like AS160, GLUT4 and glucose. AS160 is an Akt substrate (of 160 kDa) that induces translocation of GLUT4 to the plasma membrane. GLUT4 is an insulin-regulated glucose transporter. Whereas in Talbot et al. (Talbot et al., 2012), they included components that are linked to Alzheimer’s, like JNK (c-Jun NH_2_-terminal kinase, connected to embryonic development and apoptosis), IKKbeta (Inhibitor kappa B kinase, related to inflammation), ERK2 (extracellular signal-regulated kinases, increases resulting from several factos like oligomeric Abeta, oxidative & inflammatory stress factors, neurotrophins in connection to IGF1-insulin-like growth factor). Both have feedback loop from mTOR (mammalian target of rapamycin) to IRS. mTORC1 functions as nutrient/energy/redox sensor and controls protein synthesis. Only Bränmark et al. included mTORC2, a protein complex that regulates cellular metabolism as well as the cytoskeleton.

Bränmark et al. analyzed their model by interchanging the following three diabetes parameters highlighted in the red dotted boxes. In the diabetes state of the model, they include the reduced concentration of insulin receptor, GLUT4, and changed feedback from mTORC1. They extended the model by including compartmental details. They further differentiated those in active state, in plasma membrane-localized state and in the internalized state via subscripts a, m, and i, respectively, highlighted in yellow box. Whereas in Talbot et al., they analyzed their data by comparing the values of IRS1, JNK, IKKbeta, and ERK to show the link between insulin and Alzheimer’s (independent of T2D). The data that can be used in both papers for comparison include the dose-response data for AKT or PKB, and phosphorylated IRS. The one for IRS might be a little hard to compare due to the small data points considered by Talbot et al. as compared to that by Bränmark et al. However, we tried to resolve it by normalizing the data in order to make the data from two different experiments comparable.

In Talbot and colleagues’ work, their focus is mainly comparing insulin signaling in normal and AD patients without T2D. They showed that in AD patients, with or without T2D, the brain itself becomes insulin resistant. They made their study of insulin signaling in AD independent of T2D. They also highlighted that brain insulin resistant in AD is associated with IGF-1 resistance, IRS-1 dysregulation and cognitive decline. Whereas in Bränmark and colleagues work, the main result of their study is that mTORC1 activation and feedback from mTORC1 to IRS1 is the most important change going from normal to diabetic signaling. In relation to this they even tried to inhibit mTORC1 for the possibility of future development of therapeutic drug.

In this study, we extended Bränmark et al. model by integrating the components proposed in Talbot et al. (see Fig. 2). The pathway is composed of insulin signaling that inhibits gluconeogenesis in the liver, through disruption of mTORC2 binding. Particular in this model is the inclusion of JNK, IKKβ and ERK2 that are hypothesized or said to be contributory factors to the development of AD.

**Fig. 2.**
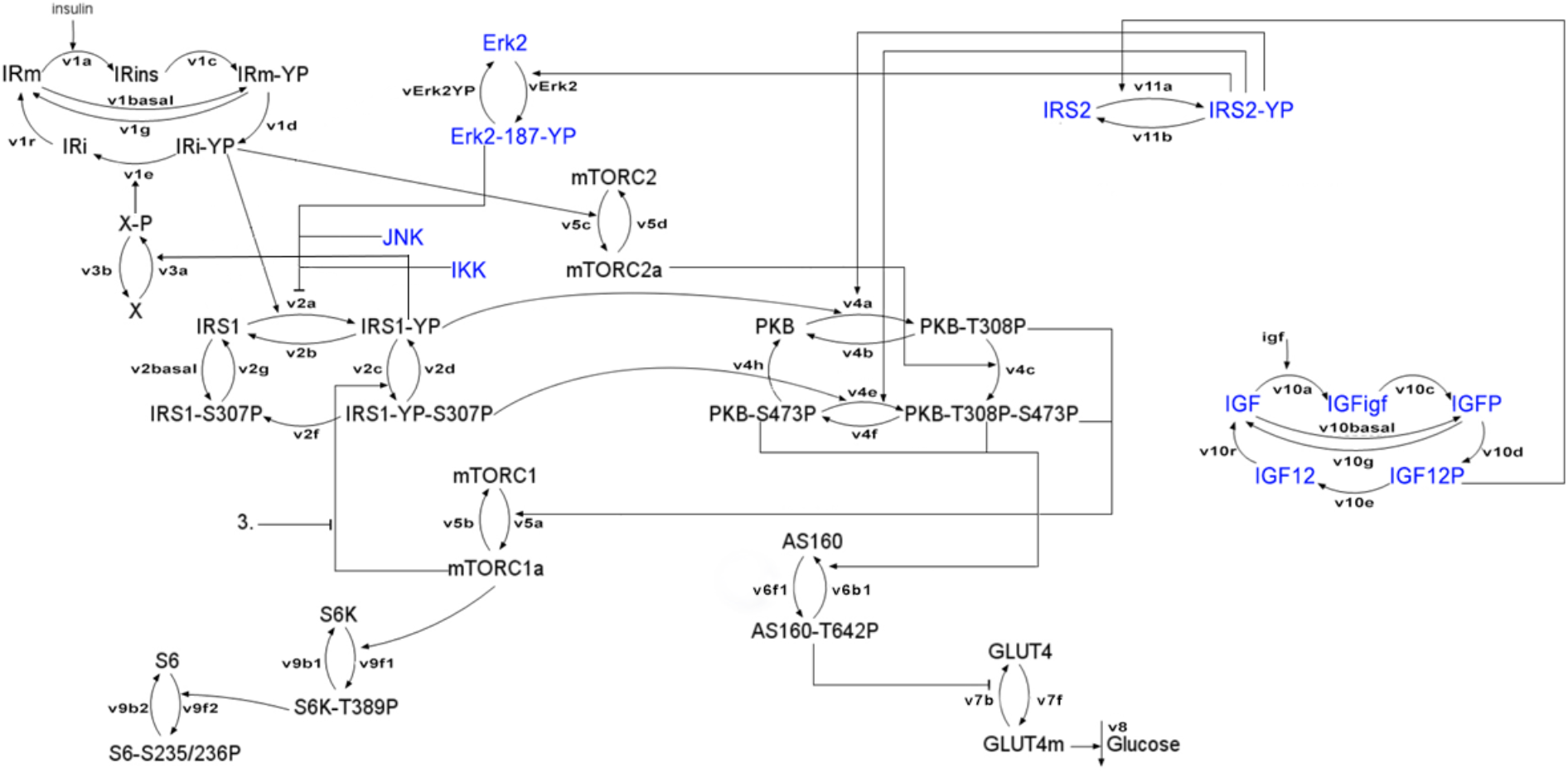
Schematic diagram of the extended model.

The set of ordinary differential equations is established based on mass action law, with consideration of hill equations. Each of the dependent variable pertains to the components/proteins/genes illustrated in the figure. Through ordinary differential equations, we are modeling the change of the concentration of these components with respect to time.

## RESULTS

As shown in Fig. 3, columns 1 and 3 show the dose responses for the indicated signaling intermediaries for non-diabetic patients (in blue dots) along with the data from Talbot et al. on the hippocampal formation of normal cases in green boxes. In contrast, columns 2 and 4 show the dose responses for the same signaling intermediaries for diabetic patients (in red dots) along with the data from Talbot et al. on the hippocampal formation of AD cases in pink boxes. The blue and red curves correspond to the simulation for non-diabetic and diabetic patients, respectively. In general, the simulated model matched Talbot’s data (those in green and pink boxes) better than Bränmark et al.

**Fig. 3.**
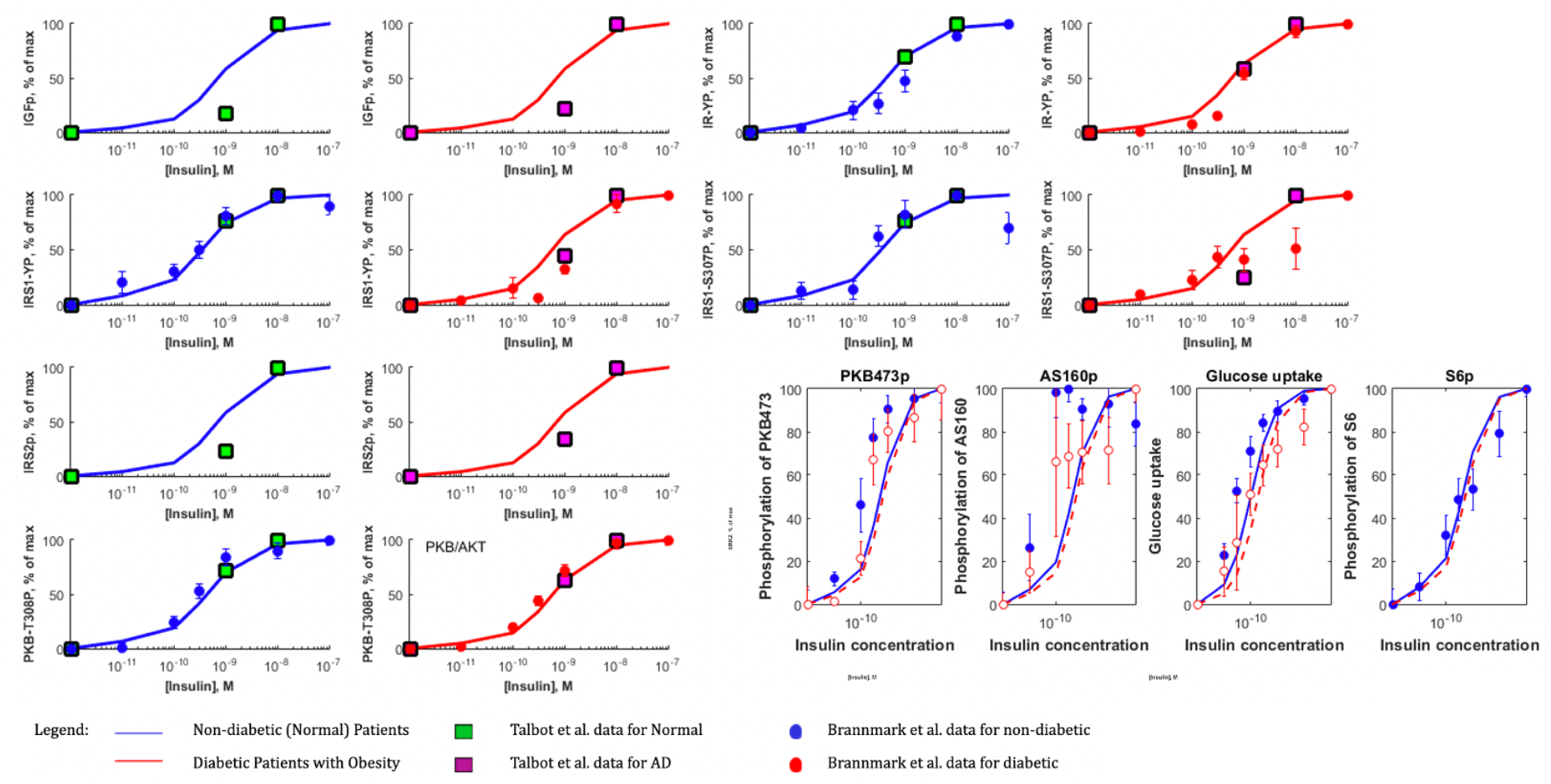
Model Simulation.

In Fig. 4, insulin is fixed at 10 nM. The simulation of PKB308p has higher concentration as compared to what Bränmark generated for plainly T2D patients. Whereas, for the simulation of PKB473p, we got lower concentration as compared to what Bränmark generated for plainly T2D patients. This may imply and suggest that for patients with T2D and AD, they will have higher PKB308p and lower PKB473p. Note that in the model (see Fig. 2), PKB or AKT is important because it inhibits GSK3beta that is associated to Amyloid Precursor Protein APP and Tau of Alzheimer’s disease. Although GSK3beta is not included in the model, but what we observed in the model simulation confirms the important role played by PKB/AKT in activating Alzheimer’s disease if T2D is not treated properly. AS160p in red dashed line matches that of Bränmark’s data for T2D patients. This indicates that fewer GLUT4 is being translocated to the plasma membrane to transport insulin-regulated glucose.

**Fig. 4.**
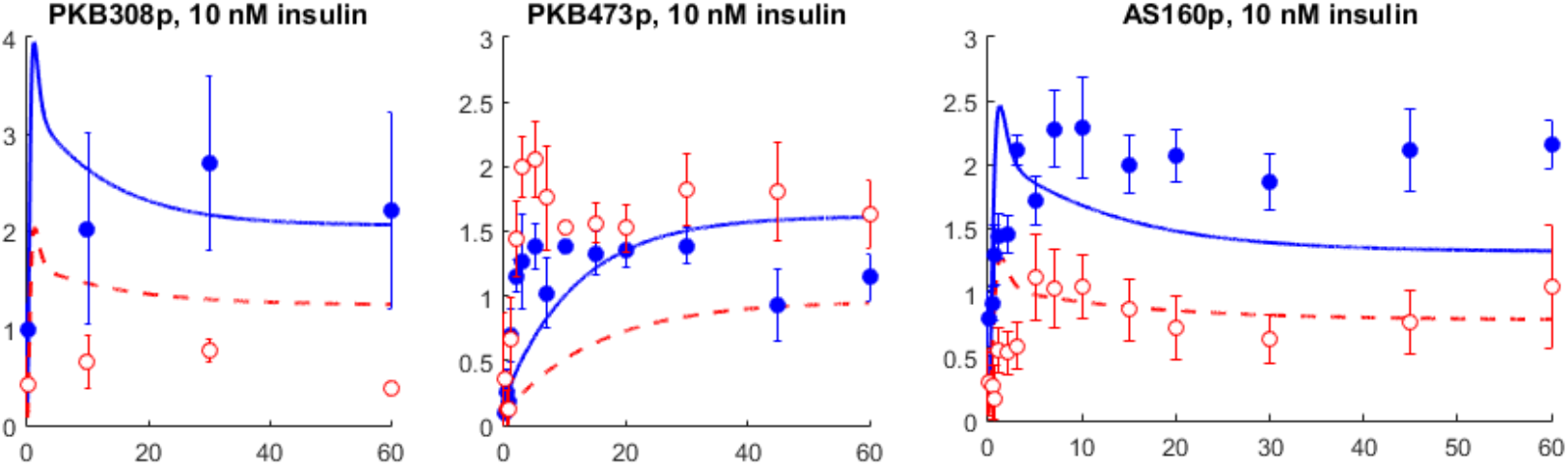
Insulin fixed at 10 nM for PKB308p, PKB473p and AS160p.

## CONCLUSION

In the simulations of the T2D-AD model, it has lower PKB473p concentration values confirming the result of Talbot in their study that brain insulin resistant in AD is associated with IGF-1 resistance. We were able to link Bränmark’sT2D relatedInsulin Resistant model and data to Talbot’s AD related insulin resistant model and data. However, due to the complexity of the model, there is still works that need to be done in the parameters used in the study.

## Supporting information

Supplementary Methodology

